# Anterior insular activity signals perceptual conflicts induced by temporal and spatial context

**DOI:** 10.1101/2024.05.29.595872

**Authors:** Katrin Reichenbach, Marcus Rothkirch, Lucca Jaeckel, Philipp Sterzer, Veith Weilnhammer

## Abstract

The signals registered by our senses are inherently ambiguous. Subjective experience, by contrast, is informative: it portrays one interpretation of the sensory environment at a time while discarding competing alternatives. This is exemplified by bistable perception, where ambiguous sensory information induces prolonged intervals of alternating unambiguous perceptual states. According to predictive-processing accounts of bistable perception, perceptual experiences in the recent past constitute a predictive context that stabilizes perception, while sensory information that is in conflict with this predictive context evokes prediction errors. These prediction errors are thought to drive spontaneous perceptual switches. We asked whether this mechanism generalizes to conflicts between other forms of predictive context and sensory information.

To this aim, we investigated the neural correlates of perceptual conflicts with temporal and spatial context during bistable perception using functional magnetic resonance imaging (fMRI). Twenty-six healthy participants viewed serial presentations of ambiguous structure-from-motion stimuli either in isolation (conflict with temporal context) or embedded in a similar but unambiguous surround stimulus (conflict with spatial context). The neural correlates of conflicts with temporal and spatial context overlapped in the anterior insula bilaterally. Model-based analyses similarly yielded common prediction error signals in the anterior insula bilaterally, right inferior frontal gyrus and right inferior parietal lobe. Together, these findings point to a generic role of these frontoparietal regions in detecting perceptual conflict and thus in the construction of unambiguous perceptual experiences.

## Introduction

Constantly faced with the challenge to resolve ambiguous sensory input, the human brain relies on prior knowledge derived from previous experiences and contextual information (see Brascamp et al., 2018 for a review). The framework of predictive processing proposes that conflicts between internal predictions (priors) and sensory input result in prediction errors, which can update the brain’s predictive model, thus minimizing prediction error in the long run (Friston, 2005; Hohwy et al., 2008). According to predictive-processing accounts of bistable perception, the spontaneous alternation between two perceptual interpretations of a single stimulus (Blake & Logothetis, 2002; Leopold et al., 1999), perceptual experiences in the recent past constitute a predictive context that stabilizes perception. Conflicting ambiguous information evokes prediction error signals and ultimately drive spontaneous perceptual switches (Brascamp et al., 2018; Hohwy et al., 2008). Neuroimaging studies have pointed to a key role of right-hemispheric frontoparietal regions including inferior frontal cortex (IFC, defined as the anterior insula bordering on the inferior frontal gyrus (IFG)) in bistable perception (see Brascamp et al., 2018 for a review). However, while neural activations in these regions have often been observed in association with spontaneous perceptual switches (Lumer et al., 1998; Sterzer & Kleinschmidt, 2007; Weilnhammer et al., 2017, 2021), their computational function has been a matter of debate (Brascamp et al., 2015, 2018; Frässle et al., 2014; Knapen et al., 2011; Zhu et al., 2022). Recent model-based fMRI has indicated that IFC detects prediction errors that are fed-forward from feature-selective regions in visual cortex (e.g., V5/hMT+ for motion), thereby signaling conflicts between the ambiguous sensory input and internal predictions derived from the temporal context of preceding perceptual states (Weilnhammer et al., 2021).

Here, we asked whether frontoparietal activity might represent a more general signature of prediction errors that extends beyond the specific case of conflict with temporal context. We devised a new experimental paradigm to test whether the processing of perceptual conflict with temporal context shares a common neural substrate with another potential source of conflict, spatial context. We used an ambiguous structure-from-motion stimulus that was presented intermittently, a technique known to stabilize perception, thus emphasizing the effect of temporal context (Leopold et al., 2002; Pearson & Brascamp, 2008). We presented the ambiguous stimulus either in isolation (single-sphere condition) or, to provide spatial context, embedded in a perceptually similar but unambiguous surrounding sphere (double-sphere condition, Figure 1).

**Figure 1.**
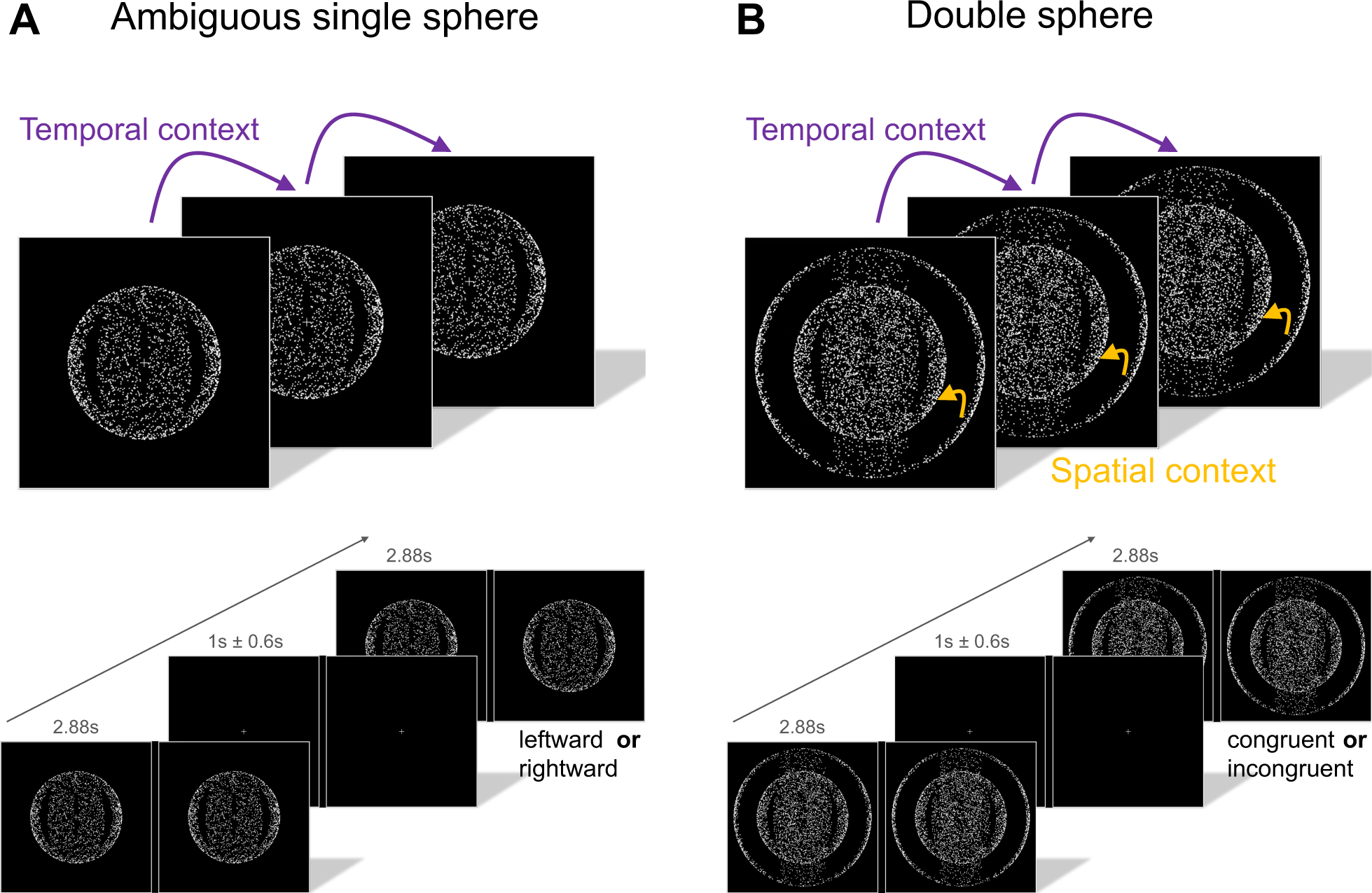
Stimulus presentation and behavioral tasks **A** Single-sphere condition. The ambiguous structure-from-motion stimulus can be perceived as a sphere of dots with a depth effect whose front surface rotates either leftward or rightward. Contextual information is provided by preceding trials (temporal context). In the single-sphere condition participants were instructed to report their percept each time the stimulus re-appeared after a fixation interval. **B** Double-sphere condition. In addition to the temporal context, spatial context is provided by the outer, unambiguous sphere. In the double-sphere condition participants were instructed to report whether both spheres moved in the same direction (congruent) or in opposite directions (incongruent).

The single-sphere condition was used to assess the neural correlates of perceptual conflict with *temporal* context. Based on the notion that switch-related neural activity reflects the peak of accumulating prediction errors (Weilnhammer et al., 2017, 2021), we contrasted trials in which participants’ perceptual reports differed from the preceding trial to those in which the perceptual report was repeated. As a neural correlate of temporal conflict, we expected to find switch-related activity in the frontoparietal cortex and V5/hMT+ as reported previously (Brascamp et al., 2018).

The double-sphere condition was designed to assess the neural correlates of perceptual conflict with *spatial* context. Based on the notion of prediction-error minimization through resolution of perceptual conflicts (see Hohwy et al., 2008) and in line with previously reported context effects in bistable perception (see Klink et al., 2012 for a review), we expected that participants’ reports would be more often congruent than incongruent with the spatial context provided by the unambiguous sphere. Critically, and under the assumption that temporal and spatial conflicts are represented by a shared neural substrate, we further expected greater neural activation in frontoparietal cortex and V5/hMT+ during percepts that were incongruent vs. congruent with the spatial context.

In addition, we devised a computational model of trial-wise perceptual prediction error, which included the preceding trial (single-sphere and double-sphere condition) and spatial contextual information (double-sphere condition) as internal predictions. The modelled prediction errors therefore signal conflicts with temporal and spatial context, which we again expected to correlate with neural activity in frontoparietal cortex and V5/hMT+.

## Materials and Methods

The study was approved by the ethics committee of Charité – Universitätsmedizin Berlin (EA1/013/20). All participants provided written informed consent and received financial compensation for their participation in the study. Exclusion criteria were MRI contraindications, neurological and/or psychiatric disorders and left-handedness. The fMRI experiment was piloted.

### Participants

Prior to the fMRI session, 58 healthy participants took part in a behavioral session outside the scanner in order to test stereopsis and to ensure a minimum amount of spontaneous perceptual switches during viewing of the intermittent ambiguous stimulus, as required for our event-related fMRI design. Behavioral testing was carried out in one or two sessions using anaglyph glasses and heterochromatic flicker photometry (n = 18) or a mirror stereoscope (n = 42). Participants were excluded based on the following perceptual performance criteria in the behavioral screening session:

1. Percentage of correct responses below 90% in the unambiguous run testing stereoscopic vision (n = 3).
2. Missed trials (no response given) exceeding 25% in any of the runs (n = 2).
3. Continuous perception of both rotation directions without clear direction (mixed percepts) in the single-sphere condition (n = 4)
4. Less than 2 perceptual switches in the single sphere run AND less than 3 switches on the inner ambiguous sphere in the double-sphere condition (n = 9). The latter criterion was added in the course of the behavioral testing and recruitment period. We noted that the first criterion was too strict when presenting stimuli intermittently, which is known to decrease perceptual switch rate (Pearson & Brascamp, 2008). Two participants having insufficient perceptual switches according to the first criterion were included at the beginning of the study due to a technical error, but fulfilled the second criterion of switches in the double-sphere condition that was added later.

Six participants were not eligible for the MRI session because of MRI exclusion criteria that were reported only after behavioral screening (n = 2) or due to drop-out from the study (n = 4).

Thus, 35 participants (the remaining 34 participants plus one additional participant who had participated in an fMRI pilot experiment) were invited to the fMRI session. One participant was excluded due to technical issues during fMRI scanning and one participant due to having difficulties to fuse the stimuli in the scanner set-up. fMRI sessions of five participants were aborted and fMRI data of two participants were excluded from further data analyses, because correct responses during fMRI scanning were below the threshold of 90%.

Thus, the final sample included 26 healthy participants (14 female, 12 male, mean age = 25.19, standard deviation (SD) = 5.73). All participants had normal or corrected-to-normal vision. The Edinburgh Handedness Inventory (Oldfield, 1971) was administered to assess handedness. All participants were right-handed, but one ambidextrous. All participants completed in-house translations of the Peters et al. Delusions Inventory (PDI, Peters et al., 1999) and the Cardiff Anomalous Perceptions Scale (CAPS, Bell et al., 2006).

If reports revealed only one percept per condition, these runs were excluded from the fMRI analysis. This was true for three participants in the single-sphere condition; and for all runs of two participants and individual runs of three participants in the double-sphere condition. One further participant was excluded from fMRI analysis of the single-sphere condition due to technical issues. One double-sphere run of another participant was excluded from behavioral and fMRI analyses because of an excessively high rate of task-unrelated button presses (42%). The average rate of task-unrelated button presses of the remaining double-sphere runs was 0.05% across participants.

### Experimental design

Visual stimuli in the fMRI setup were presented using MATLAB software (version R2020b, The MathWorks Inc.) and the Psychophysics Toolbox Version 3 (Brainard, 1997; Kleiner et al., 2007; Pelli, 1997) on an LCD monitor (NordicNeuroLab, resolution of 1024 x 768 pixels and refresh rate of 60 Hz) at the end of the scanner bore via a mirror system. Viewing distance was approximately 158 cm. We used a structure-from-motion stimulus in either ambiguous, unambiguous or combined condition (Figure 1). The sphere of dots with depth effect was built from two intersecting bands as originally introduced by Pastukhov et al. (2012) and described in Weilnhammer et al. (2021). For dichoptic stimulus presentation, we used an MRI-compatible system including a black cardboard divider and prism glasses (Schurger, 2009).

The sphere of white dots induces rotation around its vertical axis (frame: 2.27° of visual angle, double sphere: 2.06° of visual angle, single sphere: 1.37° of visual angle, individual dot size 0.03° of visual angle) and can be perceived as rotating either leftward or rightward (front surface). It is surrounded by a white frame on a uniform black background and includes a central fixation cross. For the double sphere configuration, an outer unambiguous sphere was added to the inner ambiguous sphere.

On each trial, the stimulus was presented for 2.88s followed by a jittered fixation interval of 1s ± 0.6s duration (Figure 1). Each experimental run included 80 trials and 153 - 161 task-relevant volumes. The fMRI session consisted of five runs in the following order: The first run served for stereopsis screening by using a simple unambiguous sphere (unambiguous condition) with balanced leftward or rightward rotation in pseudorandom order. Only behavioral data of this run were used in the analyses reported. In case of low correct response rate and/or reported difficulties in binocular fusion, this first run was repeated. In the second run, the single ambiguous sphere was presented (single-sphere condition). In the following three runs, the double sphere consisting of an inner ambiguous sphere and an outer unambiguous sphere representing contextual information was presented (double-sphere condition). The direction of rotation of the outer sphere was randomized, with 50% of stimuli rotating toward the right. Direction changed in average intervals of 1.94 trials (SD = 0.42).

Due to a technical issue, the randomized order of stimulus presentation across runs was identical for 11 participants. To rule out any potential confound, we included the randomization pattern type as factor in the behavioral analyses. For the sake of simplicity, randomization type was divided into two groups (group A with complete identical randomization order (n = 11) and the remaining group B (n = 15)). Two participants having the same randomization order, but different to group A, were assigned to group B. When performing an additional control analysis and excluding these two participants from mixed model analyses, results remained qualitatively identical.

Participants were instructed to report their percept (right or left in the single-sphere condition and incongruent or congruent in the double-sphere condition) via button press each time the stimulus re-appeared after the fixation interval. Responses were given on an MRI compatible button box via presses of the right hand (single-sphere condition: index finger for leftward direction and pinky finger for rightward direction and double-sphere condition: middle finger for congruent and ring finger for incongruent percept). One participant used inversed buttons in the double-sphere condition and the behavioral data were transformed accordingly.

### fMRI data acquisition

Imaging was performed on a 3 Tesla MR scanner (Siemens Magnetom Prisma) using a 20-channel head coil. Structural T1-weighted 3D magnetization-prepared rapid gradient echo (MPRAGE) images were acquired with the following parameters: 176 slices, voxel size 1 mm isotropic, TR 2530 ms, TE 4.94 ms, TI 1100 ms, flip angle 7°, FoV 256 mm.

T2-weighted 2D echo planar-imaging (EPI) images were acquired with the following parameters: 32 slices, voxel size: 3 mm x 3 mm, TR 2000ms, TE 30ms, flip angle 78°, FoV 192mm, slice thickness 3mm, slice gap 25%. Before functional scanning, a fieldmap was acquired.

### fMRI data preprocessing

Preprocessing and image analysis were performed using the SPM12 software (Wellcome Centre for Human Neuroimaging, https://www.fil.ion.ucl.ac.uk/spm/) in MATLAB (version R2019b and R2021a, The MathWorks Inc.). Preprocessing included slice-time correction with reference to the middle slice, realignment and unwarping with distortion correction using voxel displacement maps, segmentation, co-registration, spatial normalization to standard MNI space and voxel size of 3 x 3 x 3 mm^3^ and spatial smoothing with a Gaussian kernel (full-width at half-maximum of 8 mm). Realignment parameters were examined for excessive head movement defined as volume-to-volume movement > 1.5mm of translation and/or > 2 degrees of rotation (no participant was excluded due to these criteria).

### Statistical analysis of behavioral data

Statistical analysis of behavioral data were carried out using R 4.1.2 (R Core Team, 2021) and the lme4 package (Bates et al., 2015). A p-value < .05 was considered statistically significant. To explore the effects of contextual information and experimental randomization type on perceptual stability in both conditions, we fit a linear mixed-effects model. The model included perceptual stability as dependent variable and spatial context (absent/present), experimental randomization type as fixed effects including interactions. Subject was included as random-intercept term. We fit the linear mixed model as follows (Wilkinson notation):

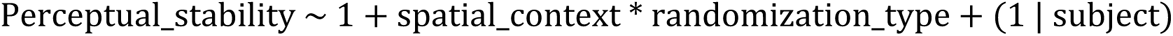

To test the hypothesis of contextual information favoring congruent perception above chance, we fit a further linear mixed-effects model. In a first step, the model included “congruency” (relative frequency of congruent percepts) as dependent variable and subject as random-intercept (Wilkinson notation):

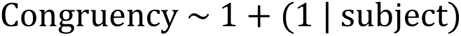

In a second step, we added session (3 runs) and experimental randomization order as fixed effects to rule out potential temporal trends or randomization design factors (Wilkinson notation):

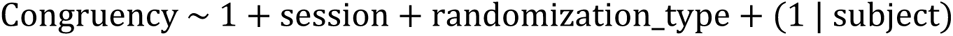

### Analysis of fMRI data

fMRI data were analyzed at two levels using the general linear model (GLM) approach in an event-related design. We estimated two different GLMs, one for the single sphere run (single-sphere condition) and one for the double sphere runs (double-sphere condition). Trials were classified based on behavioral responses (single-sphere condition: switch or no-switch derived from reported motion directions and double-sphere condition: incongruent or congruent). At the first level, we defined two regressors of interest (single-sphere condition: switch and no-switch; double-sphere condition: incongruent and congruent). Missed and multiple or task-unrelated button presses and unclear perceptual switch states (first trial, trials following a trial with no button press) in the single-sphere condition as well as the head motion parameters were included as nuisance regressors. All regressors (except head motion parameters) were modelled as stick functions aligned to stimulus onset with a duration of zero seconds and convolved with the hemodynamic response function. A high-pass filter with a cut-off at 128 seconds and an autoregressive model of order one were applied.

Individual t-contrast maps were created for each participant by subtracting no-switch trials from switch trials (switch > no-switch) for the single-sphere condition and by subtracting congruent trials from incongruent trials (incongruent > congruent) for the double-sphere condition, respectively.

### Computational modeling approach

Next, we used computational modeling to test the representation of perceptual prediction across the cortex. We applied a generative model based on a generalized linear mixed model to predict trial-wise perceptual states y(t) that indicate whether the front surface of the ambiguous sphere was perceived as rotating toward the left (y(t) = 0) or toward the right (y(t) = 1). We computed weights including two sources of influence on perception: preceding perceptual states (*temporal*, single-sphere and double-sphere condition) and spatial context information (*spatial*, double-sphere condition). In the single-sphere condition, perceptual states were only predicted by the preceding perceptual state y(t - 1) and an intercept α:

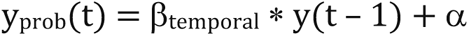

In the double-sphere condition, perceptual states were predicted by the direction of rotation of the contextual sphere s(t) and the preceding perceptual states y(t - 1). Please note that in the double-sphere condition, perceptual states y(t - 1) were inferred indirectly through the congruency response of the participants. β_temporal_ * y(t – 1) therefore represented the temporal context and β_spatial_ * s(t) the spatial context:

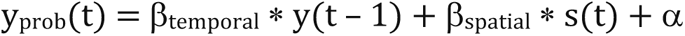

Parameters were estimated using independent linear mixed effects models with random intercepts and slopes per participant (R package lme4). We computed trial-wise prediction errors via the absolute difference between the predicted and reported perceptual states:

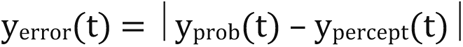

Further, we used the estimated prediction error values from the model for statistical analysis of fMRI data. We set up two GLMs (single-sphere and double-sphere condition, respectively) in SPM12 each containing the onsets of all trials as stick regressors and added y_error_ as a parametric regressor. We mean-centered the parametric regressor over condition and did not apply orthogonalization of parametric modulations in SPM12. We computed individual contrast images (y_error_ > baseline) at the single subject level.

### Region of interest definition and analysis

With respect to our hypothesis on prediction error related activity during bistable perception, we focused on five ROIs defined by literature-based coordinates (Weilnhammer et al., 2021): left anterior insula (x, y, z) = (-30, 23, -1), right anterior insula (x, y, z) = (33, 23, -1), right IFG (x, y, z) = (45, 17, 23), right inferior parietal lobule (IPL) (x, y, z) = (39, -37, 50), right V5/MT (x, y, z) = (45, -64, 8). Using MarsBaR toolbox (Brett et al., 2002), we created spheres of 6 mm radius based on the above coordinates and intersected them to anatomical masks (left insula, right insula, right inferior parietal gyrus and the combination of pars opercularis and pars triangularis of the right inferior frontal gyrus) taken from the AAL ROI library (Tzourio-Mazoyer et al., 2002). As the anatomical location of V5/hMT+ is known to vary among individuals (Bridge et al., 2014), the ROI of V5/hMT+ was not intersected to an anatomical mask and centered solely on the above coordinates. Further, we used the MarsBaR toolbox (Brett et al., 2002) to extract mean parameter estimates for the five ROIs in each individual contrast in the conventional and model-based analyses, respectively. These estimates were further analyzed in R software (R Core Team, 2021) using two-sided one-sample t-tests and Bonferroni correction for the number of ROIs (n = 5).

### Whole-brain analyses

In addition to the ROI-based analyses, we performed whole-brain analyses. We submitted the individual contrast images to a one-sample t-test at the second level and applied a familywise error (FWE)–corrected significance level of p = .05. We used AAL3 (Tzourio-Mazoyer et al., 2002) for anatomic labeling and MRIcroGL (Rorden & Brett, 2000) for visualization of brain imaging data.

## Results

### Behavioral results

Correct responses in the Unambiguous condition were at 98.82 % (SD = 2.7). Mean percentage of missed trials (no button pressed) over participants was 1.14 % (SD = 1.29). Perceptual stability was defined as the relative frequency of no-switch trials (e.g. a perceptual stability of 100% indicates that no switches occurred). In the single-sphere condition, perceptual stability across participants was 87.59 % (SD = 15.4) and the mean absolute number of perceptual switches was 9.15 (SD = 10.84, range 0-35) (Figure 2A). Perceptual stability of the inner ambiguous sphere in the double-sphere condition was 60.09 % (SD = 12.08). In the double-sphere condition, the mean of congruent percepts was 77.06 % (SD = 19.61) (Figure 2B).

**Figure 2.**
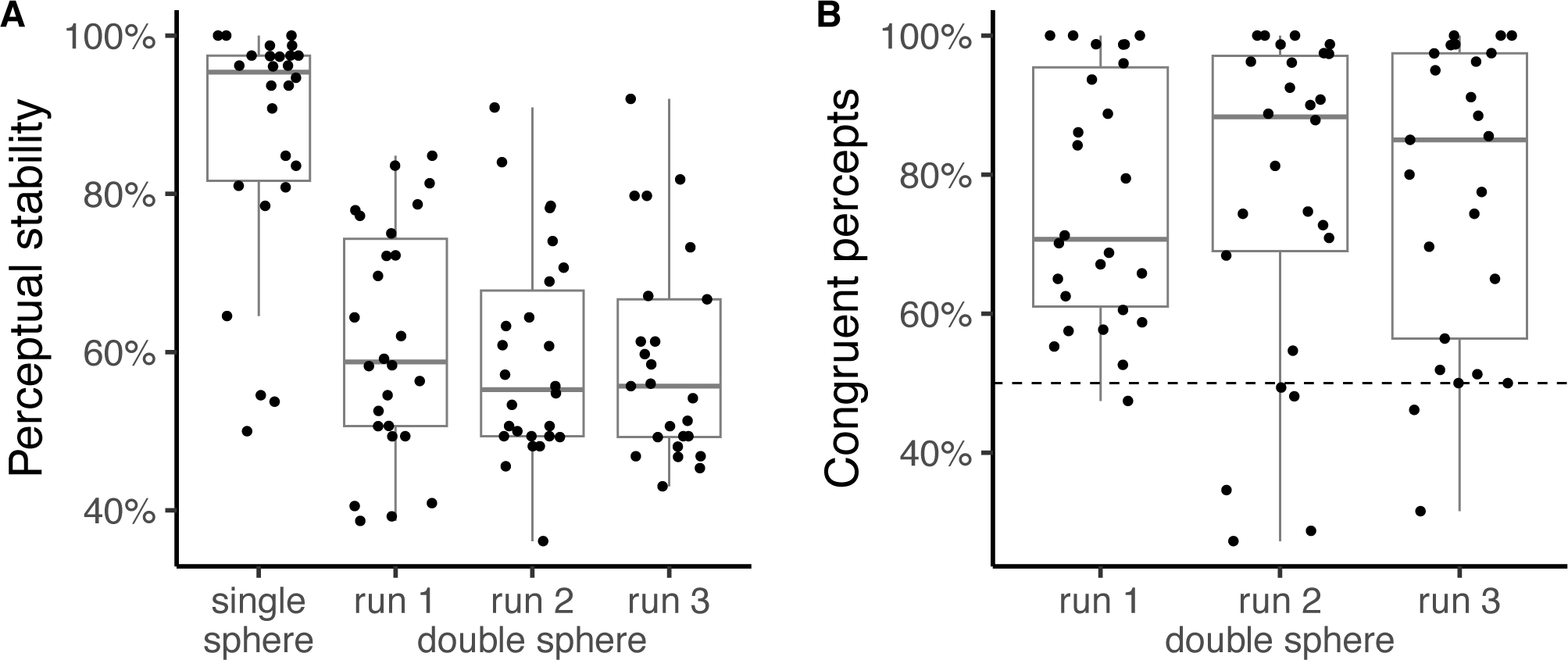
**A** Perceptual stability distributions in the single-sphere (no context) and double-sphere condition. Perceptual stability denotes the relative frequency of no-switch trials. Thus, the higher the perceptual stability the less perceptual switches occurred. Boxplots are combined with jittered data points for each participant. Boxplots visualize the median, the 25^th^ percentile (first quartile) and the 75^th^ percentile (third quartile) of the data at the middle, lower and upper hinges. Vertical whiskers indicate 1.5 fold inter-quartile ranges of the data. **B** Distribution of congruent percepts in the double-sphere condition. Congruency indicates perception of the same motion direction of the rotating inner ambiguous and the outer unambiguous sphere. Congruency denotes the relative frequency of surround-congruent percepts. Boxplots combined with jittered data points for each participant and run. The dashed line at 50% denotes chance level.

Response times in the single-sphere condition did not differ between reported left (mean = 1.27 s, SD = 0.42) or right motion direction (mean = 1.34 s, SD = 0.44; p = .62, Wilcoxon signed-rank test; Shapiro–Wilk normality test indicating non-normal distribution, W = .94, p = .02). Response times in the double-sphere condition yielded a significant difference between incongruent (mean = 1.69 s, SD = 0.34) and congruent percepts (mean = 1.37 s, SD = 0.29; t_(23)_ = 5.84, p < .001, paired t-test; Shapiro–Wilk normality test indicating normal distribution, p = .51).

There was a trend towards a global bias towards one of the two possible percepts (left motion direction) across participants in the single-sphere condition (p = .075, Wilcoxon signed-rank test against chance level; Shapiro–Wilk normality test indicating non-normal distribution, W = .91, p = .026).

The linear mixed-effects model showed a significant effect of spatial context (beta estimate = -0.231, standard error (SE) = 0.041, p < .001), indicating a decrease of perceptual stability for stimuli presented with contextual information (double-sphere condition). This was expected under the hypothesis of a significant effect of spatial context on perception, since the direction of rotation of the unambiguous outer sphere changed randomly across trials. We observed no effect of experimental randomization order (beta estimate = 0.056, SE = 0.057, p = .33). We did not find interaction effects between context and randomization (beta estimate = -0.078, SE = 0.054, p = .15).

Further, we tested congruency with spatial context against chance level. This indicated a significant positive effect of spatial context on the perception of the ambiguous inner sphere (beta estimate = 0.771, SE = 0.038, p < .001) and (t_(25.0493)_ = 7.03, p < .001, two-sided t-test against 50%). We did not find any significant effect of session or randomization type on congruency (beta estimate = 0.037, SE = 0.079, p = .64).

### fMRI results

#### Conventional fMRI analysis

As expected on the basis of previous work (Weilnhammer et al., 2017, 2021), ROI analysis of the single-sphere condition for the contrast switch > no-switch revealed significant increases in activity in left and right anterior insula, right IFG and right IPL ROIs during switch trials (Figure 3 and 4A and Table 1). We did not find greater activation for switch trials compared to no-switch trials for the right V5/hMT+ ROI. Analysis of the double-sphere condition for the contrast incongruent > congruent showed significant increases in activity in left and right anterior insula and right V5/hMT+ ROIs as well as a trend-wise significant effect in the right IFG (Figure 3 and 4B and Table 1). We did not find greater activation for surround-incongruent trials compared to surround-congruent trials in the right IPL ROI. The findings in both conditions indicate a common neural correlate of perceptual conflicts in the anterior insula bilaterally, while the other hypothesized regions showed signification activations in only one of the conditions.

**Figure 3.**
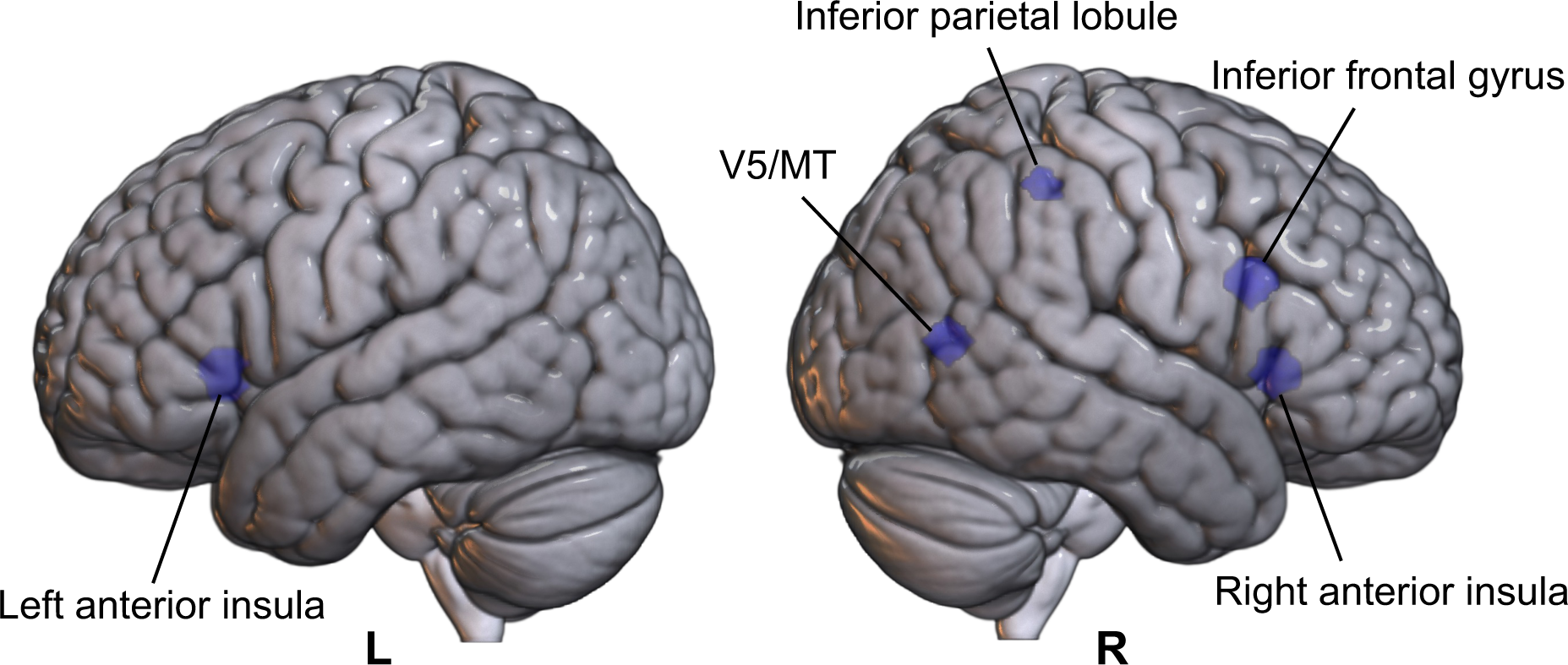
Schematic illustration of the regions of interest (ROI) overlaid on rendered MNI152 brain templates.

**Figure 4.**
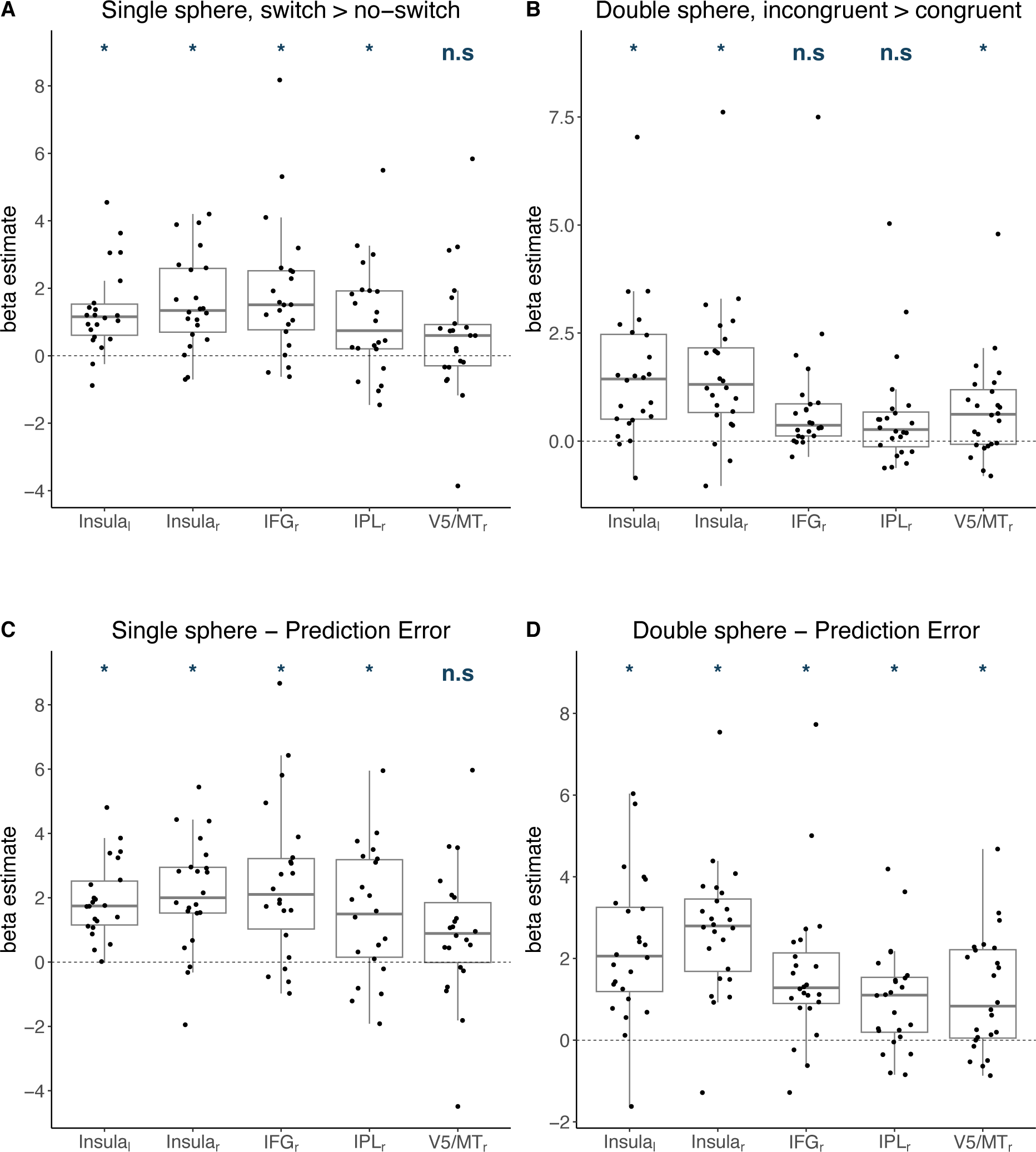
**A, B** Conventional ROI analysis. Beta estimates of brain activity for the contrasts switch > no-switch (single-sphere condition, A) and incongruent > congruent (double-sphere condition, B) in five hypothesized regions of interest: left anterior insula, right anterior insula, right inferior frontal gyrus, right inferior parietal lobule and V5/MT. **C, D** Prediction-error model-based ROI analysis. Beta estimates derived from the modulation of fMRI activity by modelled prediction errors. Boxplots are combined with jittered data points for each participant. Asterisks denote significant differences from zero (*p < .05). Abbreviation: n.s., not significant.

**Table 1.**
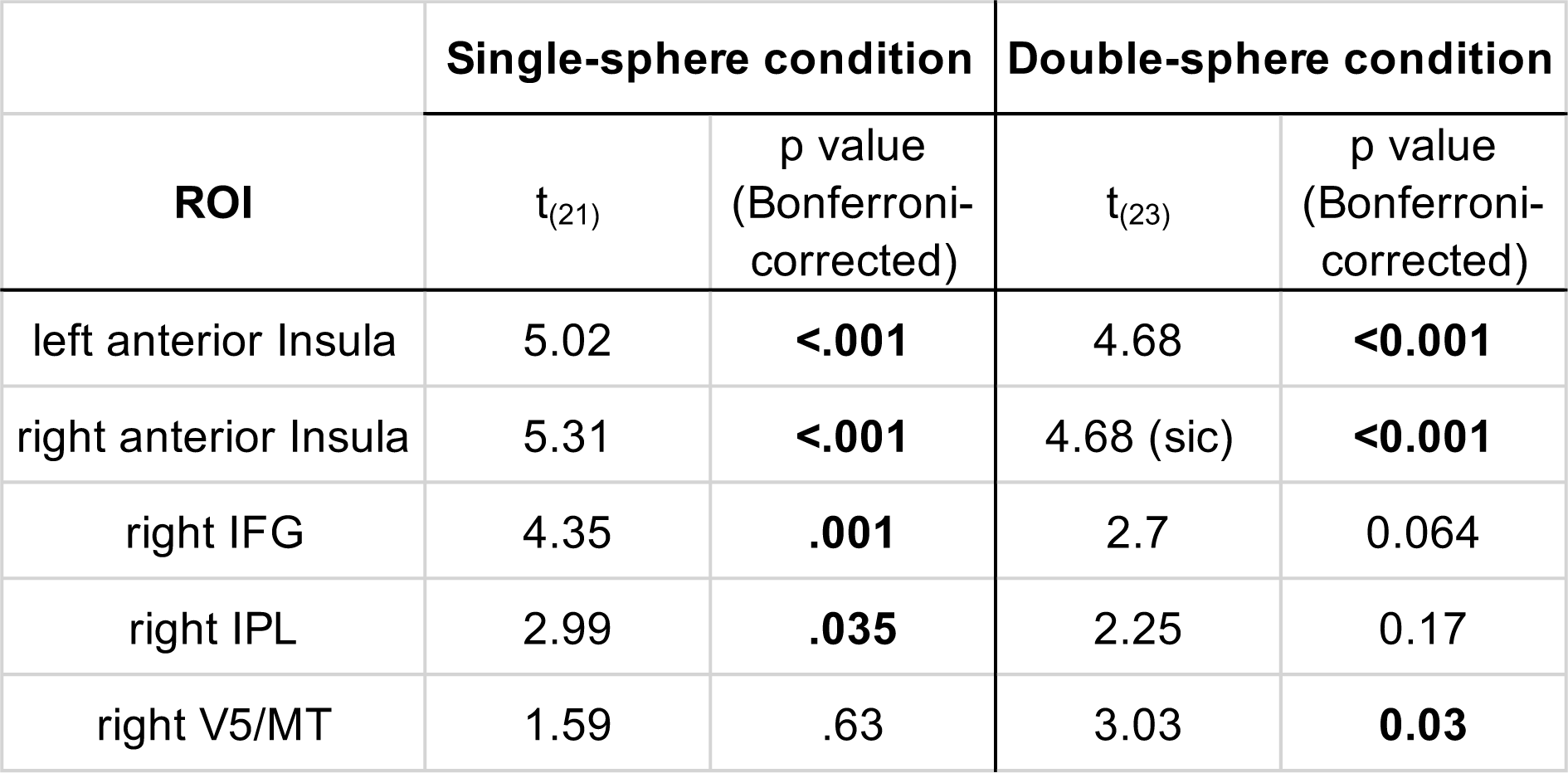
Conventional ROI analysis. Beta estimates for both contrasts (switch > no-switch and incongruent > congruent) using one-sample t-tests.

Given that the perception of the stimuli varied largely between participants (see extended Data Figure 2-1), we performed additional control analyses and tested whether the number of events per condition (switches and incongruent trials, respectively) might have influenced the magnitude of beta estimates of each ROI (extended Data Figure 4A,B-1). We found correlations between the frequency of events and activation strength in the anterior insula bilaterally and the right IFG. After excluding 1) participants with less than three switches in the single-sphere condition (remaining n = 15) and 2) all runs with less than 10% incongruent trials in the double-sphere condition (remaining n = 19), ROI effects remained robust in the right frontal area (extended Data Table 1-1). Further, additional anatomical ROI analyses (extended Data Figure 4A,B-2) did not reveal significant correlations in other brain regions.

Additional whole-brain analysis of switch-related activity in the single-sphere condition (contrast switch > no-switch) did not reveal any surviving clusters after FWE correction at a significance level of p < .05. Whole-brain analysis of incongruent activity in the double-sphere condition (contrast incongruent > congruent) showed activations in the supplementary motor area (extended Data Table 1-2).

#### fMRI Prediction Error Model

To test our hypothesis of prediction error signaling, we examined the modulation of functional fMRI activity during visual onset by prediction error estimates derived from our computational modelling approach using a generalized linear mixed model to predict trial-wise perceptual states. As hypothesized, ROI analyses revealed correlations of neural activity with perceptual prediction errors in the anterior insula bilaterally, right IFG and IPL for both the single- and the double sphere condition (Figure 3, 4C, 4D and Table 2). Neural activity in the V5/hMT+ region correlated with prediction errors in the double-sphere but not in the single-sphere condition.

**Table 2.** ROI analysis of Prediction Errors. Beta estimates for both conditions using one-sample t-tests.

| ***Neural coding of prediction errors*** | **Single-sphere condition** |  | **Double-sphere condition** |  |
| --- | --- | --- | --- | --- |
| **ROI** | $t_{(21)}$ | p value<br>(Bonferroni-corrected) | $t_{(23)}$ | p value<br>(Bonferroni-corrected) |
| left anterior Insula | 7.58 | **<0.001** | 6.28 | **<0.001** |
| right anterior Insula | 5.75 | **<0.001** | 8.17 | **<0.001** |
| right IFG | 4.79 | **<0.001** | 4.43 | **0.001** |
| right IPL | 3.53 | **0.01** | 3.95 | **0.003** |
| right V5/MT | 2.05 | 0.27 | 3.92 | **0.003** |

Additional whole brain analyses showed significant correlation with activity in the left insula in the single-sphere condition (Figure 5 and extended Data Figure 5-1); and in the insula bilaterally, left superior frontal gyrus, right IFG and left precentral gyrus, among other regions in the double-sphere condition (Figure 5 and extended Data Figure 5-2).

**Figure 5.**
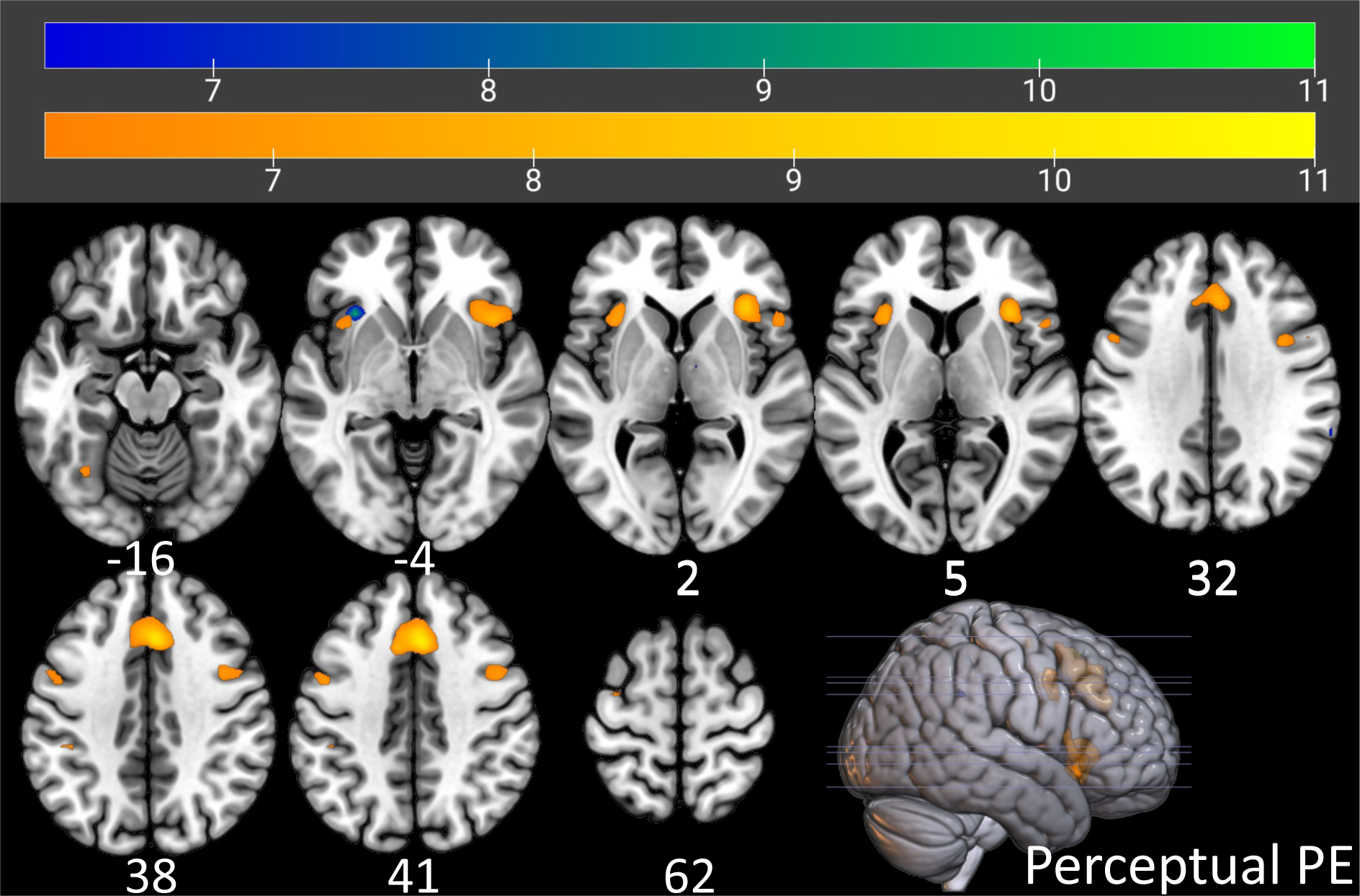
Prediction error-related brain activity in both the single-sphere and double-sphere condition. Smoothed whole-brain maps (p < .05, FWE-corrected) of perceptual prediction errors are overlaid on an SPM152 MNI brain template using MRIcroGL. Color bars indicate t values (Single-sphere condition: blue green colors, double-sphere condition: warm colors). Numbers below axial slices indicate z-coordinates.

## Discussion

This study investigated the neural correlates of conflict during perceptual inference. We used serial presentations of an ambiguous stimulus that was shown either in isolation or within an unambiguous surround. We thereby created conflicts with both temporal and spatial context, which we modelled as prediction errors in a predictive processing framework. In line with previous findings (reviewed in Klink et al., 2012), our behavioral results confirm that spatiotemporal context influences perceptual inference. At the neural level, our findings provide support for an overlapping neural substrate of perceptual conflicts with temporal and spatial context: Activity in bilateral anterior insula did not only correlate with conflict-related percepts that violated temporal and spatial context in the conventional analysis, but also with modelled prediction errors that integrated temporal and spatial context with ambiguous sensory data into a common decision variable.

At the behavioral level, we found a strong effect of temporal context on perception as evidenced by above-chance trial-to-trial perceptual stability in the single-sphere condition, in line with previous work using intermittent presentation of ambiguous stimuli (reviewed in Pearson & Brascamp, 2008). In the double-sphere condition, we observed a significant effect of spatial context, as indicated by the above-chance congruency between perception of the ambiguous stimulus and the unambiguous spatial context. Congruent perception came along with a reduced stability of the ambiguous sphere, thus resulting in an increased perceptual switch rate compared to the single-sphere condition. Our finding that surround-congruent percepts outweighed surround-incongruent percepts is consistent with previous studies (for a review see Klink et al., 2012). For example, Sundareswara & Schrater (2008) employed the Necker cube which was flanked with unambiguous cubes and showed that context-consistent interpretations were perceptually dominant. In sum, our findings align with the predictive processing premise that predictions induced by temporal and spatial context contribute to the resolution of perceptual ambiguity.

At the neural level, we asked whether frontoparietal cortex plays a general role in prediction error signaling during perceptual inference. Previous findings indicate an involvement of frontoparietal regions in switches during bistable perception and binocular rivalry (reviewed in Brascamp et al., 2018), explaining them via hierarchical models (Kanai et al., 2011; Megumi et al., 2015; Weilnhammer et al., 2017, 2021). Based on predictive processing, we interpreted switch-related neural activations as the endpoint of a dynamic accumulation of prediction errors evoked by conflict with temporal context and asked whether this would generalize to conflicts with spatiotemporal context.

Conventional analyses revealed bilateral anterior insular cortex as a common neural substrate of perceptual conflicts with temporal and spatial conflict. In addition, we found significant activation in the right IFG and IPL ROIs in the in the single-sphere but not in the double-sphere condition.

The model-based analyses confirmed bilateral anterior insula as a neural correlate of prediction errors across conditions. The same was true for right IFG and right IPL. Of note, previous work has suggested that prediction errors do not originate in IFC, but are fed-forward from feature-selective regions of visual cortex, such as V5/hMT+ for structure-from-motion stimuli (Megumi et al., 2015; Weilnhammer et al., 2013, 2017, 2021). Here, we found enhanced activity in the V5/hMT+ ROI for conflicts with spatiotemporal context (double sphere), but not with temporal conflict alone (single sphere) in both analyses. We speculate that this inconsistency with previous work and the partially divergent results in both conditions may result from the lack of statistical power in the single-sphere condition.

Taken together, our fMRI results indicate that conflicts with temporal and spatial context lead to activity in bilateral anterior insular cortex. This suggests that anterior insular activity signals prediction errors induced by conflicts with various sources of contextual information (Klink et al., 2012). Our findings support the hypothesis that the anterior insula registers prediction errors arising from perceptual conflicts, thus playing a key role in the construction of unambiguous perceptual states from ambiguous sensory information through prediction error minimization (Dwarakanath et al., 2023; Weilnhammer et al., 2021). This aligns with previous findings that highlight the insula as a domain-general locus of perceptual, cognitive and reward prediction errors (see Corlett et al., 2022 for a review).

Importantly, the correlation of anterior insular activity with temporal and spatial conflicts must be interpreted in the light of the multifunctional and complex role of the insula. The function of the insular region has been associated, among other functions, with salience processing and executive functioning (Molnar-Szakacs & Uddin, 2022; Uddin, 2015). As reviewed by Sterzer & Kleinschmidt (2010), activation of anterior insula is a widespread phenomenon apparent in a multitude of perceptual tasks, possibly mediating sensory alertness driven by salient events or challenging task demands.

Our supplemental control analyses of conventional contrast weights showed that activity in the anterior insulae and right IFG correlated negatively with the frequency of events in the single-sphere (switches) and double-sphere condition (surround-incongruent percepts). This makes sense from the perspective of predictive processing, as rare events occur in the context of strong predictions, which lead to larger prediction errors when they are violated. One concern, however, may be that rare events have effects at the level of arousal, salience, or task-related behavior that may also explain increases in cortical activation. Likewise, rare events may lead to overestimated betas as statistical artifacts. However, the anterior insular effects remained robust after excluding participants with a limited number of events per condition, rendering such an explanation unlikely. Moreover, when assessing the correlation of the frequency of perceptual events with fMRI activity in a larger set of anatomical ROIs, we did not find a general pattern of negative correlations between the respective conventional beta-estimates and the number of events (see Extended Data).

Several limitations need to be considered. Since we did not study the processing of temporal and spatial context in the absence of report, we cannot rule out a contribution of the participants’ execution of motor reports to our fMRI results. However, our task design required a motor response at each stimulus presentation and regardless of the perceptual outcome. As we used differential contrasts for each condition (switch > no-switch in the single-sphere condition and incongruent > congruent in the double-sphere condition) in the conventional fMRI analysis, neural effects of motor responses were likely neutralized. Notwithstanding, future research using no-report paradigms on bistable perception is required (Brascamp et al., 2018; Whyte et al., 2022). Furthermore, studies with larger sample sizes are needed to replicate our findings, as we had to deal with a limited and unequal number of events and a small sample size. Further, intermittent stimulus presentation is but one technique of percept stabilization, potentially involving perceptual memory processes (for a review see Pearson & Brascamp, 2008) and additional activity in frontoparietal and temporal regions (Sterzer & Rees, 2008; Wang et al., 2013). Our model-based findings suggest that the bilateral anterior insulae extending into the right IFG are involved in signaling prediction errors elicited by intermittently presented structure-from-motion stimuli. It remains an important task for future research to determine whether spatiotemporal conflicts lead to similar patterns of prediction error signaling during continuous stimulus presentation and in other types of bistability, such as binocular rivalry, figure-ground illusions, static depth inversion images such as the Necker Cube, or apparent motion.

In conclusion, our findings point to a general representation of perceptual prediction errors in bilateral anterior insular cortex. According to predictive processing, temporal and spatial context provide internal predictions that contribute to the resolution of perceptual ambiguities and conflicts. Of note, changes in the precision of internal predictions have repeatedly been linked to psychotic symptoms such as hallucinations or delusions (see Adams et al., 2013; Petrovic & Sterzer, 2023; Sterzer et al., 2018 for reviews). Further work studying context effects in bistable perception are therefore warranted, as a deeper understanding of their neurocomputational correlates might help us to better understand alterations of perception in neuropsychiatric disorders such as schizophrenia (see Teufel & Fletcher, 2020 for a review).

